# Brain-Penetrant NF-κB and NLRP3 Targeting Nanoligomers are Therapeutic in Amyotrophic Lateral Sclerosis (ALS) and Alzheimer’s Disease (AD) Human Organoid and Mouse Models

**DOI:** 10.1101/2024.03.07.583991

**Authors:** Sadhana Sharma, Devin Wahl, Sydney Risen, Vincenzo S. Gilberto, Anushree Chatterjee, Julie A. Moreno, Thomas J. LaRocca, Prashant Nagpal

## Abstract

Millions of people suffer worldwide from neurodegenerative diseases ranging from rapidly progressing and fatal motor neuron diseases like Amyotrophic Lateral Sclerosis (ALS) to more chronic illnesses such as frontotemporal dementia (FTD) and Alzheimer’s disease (AD). A growing number of studies have implicated neuroinflammation as a key and causative phenomenon and an important target for novel therapeutics for these diseases. Neuroinflammation is characterized by reactive glial cells that produce pro-inflammatory neurotoxic cytokines. Our previous studies have shown a brain-penetrant Nanoligomer cocktail (NI112) inhibiting the neuroinflammation mediators nuclear factor kappa-light-chain-enhancer of activated B cells (NF-κB) and NOD-like receptor family, pyrin domain containing 3 (NLRP3) is a safe, targeted, and effective neurotherapeutic drug. Here, we show that a four-week NI112 treatment is therapeutic using: 1) an ALS-FTD 3D human motor neuron organoid model of tar DNA binding protein 43 (TDP-43, a key contributor to ALS pathology) overexpression (knock-in); 2) an AD model of APOE4/APOE4 (AD risk allele) double mutation in human neurons comprising a 3D human prefrontal cortex (PFC) organoid; and 3) multiple *in vivo* (mouse models) of the same/related conditions. In 3D organoids made from healthy motor neurons (HMN negative control) and TDP-43 overexpressing (or ALS organoids), we monitored the mean firing rate using calcium signaling as a functional output, while measuring TDP-43 and other key neurodegeneration biomarkers. After 4 weeks, we observed a massive improvement in the mean firing rate of NI112-treated ALS organoids compared to untreated ALS organoids, which was more comparable to healthy HMN organoids. Similarly, we found a significant decrease in neurodegeneration markers like amyloid beta 42 (Aβ42) in NI112-treated AD organoids compared to untreated AD organoids (Aβ42 comparable to healthy PFC organoids). In the mouse ALS (SOD1-G93A) model, we observed behavioral improvements and restoration of motor function (e.g., grip strength) in NI112-treated mice, and in mouse AD model mice (radiation-induced accelerated neuropathology in APP/PS1, and rTg4510 phospho-tau), we observed improved cognition. In both models, we also found an accompanying reduction in neuroinflammation and reduced neuropathology. These results show the promise for further testing and development of neuroinflammation-targeting Nanoligomers to benefit patients suffering from debilitating neurodegenerative diseases like ALS, FTD, and AD.

## INTRODUCTION

### Amyotrophic Lateral Sclerosis (ALS) is a fatal motor neuron disease with no cure

ALS is classified as a “rare” neurodegenerative disease affecting ∼30,000 people in the US.^1^ ALS is progressively neurodegenerative, and patients lose the ability to walk, talk, eat, and eventually breathe. The disease is fatal with an average lifespan of 2-5 years, with no known cures, and the majority of cases (∼90%) have no genetic cause or family history. A growing number of studies have now implicated neuroinflammation as a key or causative mechanism in ALS pathology,^2–4^ and therefore a target for new therapies. While pan-immunosuppressive corticosteroids are widely available, poor clinical results with a high risk of side-effects^5^ indicate a more targeted approach is necessary to slow disease progression or achieve remission.

### Neuroinflammation-mediated pathology causes ALS disease progression

Several inflammatory cellular pathways are implicated in ALS pathophysiology. For example, activated microglia have been shown to switch from the M2 to M1 phenotype and express a neurotoxic cocktail of pro-inflammatory cytokines and reactive oxygen species (ROS) in ALS disease progression. ^2,3^ Astrocytic expression of mutated superoxide dismutase 1 (mSOD1) has also been directly linked to motor neuron pathology and ALS disease phenotype.^4^ Furthermore, CD4+ T cell infiltration has been associated with ALS progression,^6,7^ and ALS patients have increased peripheral monocytes and macrophage invasion in the spinal cord, which contributes to neuropathology and motor neuron toxicity.^8^ Infiltrating immune cells release pro-inflammatory cytokines, such as tumor necrosis factor-alpha (TNF-α), Interleukin-1beta (IL-1β), and Interleukin-6 (IL-6), which actively contribute to inflammation, tissue damage, and loss of motor neurons within the brain and spinal cord. Decreases in endogenous T regulatory (Treg) cell levels have also been linked to ALS progression and accompanying loss of neuroprotective M2 microglia.^9^ TDP-43 is another key risk neuropathological factor for ALS, AD,^10–13^ as well as many other neurodegenerative diseases. While previous studies focused on TDP-43 transfer between neurons, its role in neuroinflammation as well as expression through reactive microglial cells and astrocytes has been shown to induce neuronal toxicity.^13^ Finally, another key autoimmune response shown to contribute to ALS pathology is the activation of the complement system. Specifically, C1q and C5a components are key immune cell recruiters, whereas C5-C9 form cytotoxic membrane attack complexes, and are found to be elevated in the spinal cords of ALS patients.^14,15^

### Neuroinflammation in Alzheimer’s disease (AD)

50 million people worldwide suffer from dementia, and ∼30 million suffer from Alzheimer’s disease (AD), the most common cause of dementia.^16,17^ Brain aging involves declines in cognitive function and pathological events that are precursors to AD (e.g., the deposition of amyloid beta (Aβ) and tau), and neuroinflammation is a central contributor to all of these events.^18–23^ Key pathological features of AD are the deposition of amyloid beta (Aβ) and hyperphosphorylated tau.^24,25^ Similar to ALS, in AD neuroinflammation is characterized by innate immune activation and the production of pro-inflammatory, CNS-toxic cytokines, especially by glial cells.^18–23^ It plays a central role in AD, as reflected by the fact that many AD risk genes have innate immune functions (e.g., CD33, TREM2).^26^ However, neuroinflammation precedes Aβ/tau pathology, increases further in response to pathology, and directly reduces cognitive function.^27,28^ As such, an important goal for pharmaceutical research is to identify interventions for reducing neuroinflammation in the context of AD-related misfolded protein pathology.

### NF-κB and NLRP3 inhibition using Nanoligomers could offer a safe and targeted approach as a potential ALS and AD therapeutic

NF-κB (nuclear factor kappa-light-chain-enhancer of activated B cells) is arguably one of the most important therapeutic targets for ALS^10^ given its direct role in inflammatory pathways outlined above,^29,30^ and links to several other ALS genes such as Chromosome 9 Open Reading Frame 72 (c9orf72),^31^ Superoxide Dismutase 1 (SOD1),^32^ TDP-43,^11^ Fused in Sarcoma (FUS),^33^ Optineurin (OPTN),^34^ and TANK-binding kinase 1 (TBK1).^35^ While NLRP3 (NOD-like receptor family, pyrin domain containing 3) inflammasome is also activated during microglial activation in ALS disease,^36,37^ NLRP3 inhibition alone does not reduce ALS inflammation and neuropathology^38,39^ further highlighting the need for targeting more relevant pathways such as NF-κB.

Among several neuroinflammation targets, NF-κB also has a key role in AD.^40^ NF-κB is a central upstream transcription factor involved in many inflammatory processes,^41^ as well as in NLRP3-associated inflammatory responses^42^ to both extracellular and intracellular stimuli,^43^ many of which are implicated in AD. NF-κB is arguably one of the most important therapeutic targets for AD^40^ given its direct role in inflammatory pathways outlined above, and links to several other AD risk factors such as APOE4,^44^ glutamate,^45^ and several micro-RNAs^46^ such as miRNA-125b,^47^ miRNA-34a,^48^, and others. For example, multiple age- and AD-related signals (e.g., systemic inflammatory processes, immune dysregulation, etc.) elicit NF-κB signaling via Tumor Necrosis Factor (TNF) receptors and Toll-Like Receptors (TLRs),^49^ and NF-κB is also activated in response to intracellular stressors that increase with aging (e.g., reactive oxygen species and DNA damage).^50,51^ NLRP3 plays a major role in many of these events, and in responses to pathogen-associated molecular patterns (e.g., LPS, viral RNA), as well as mitochondrial reactive oxygen species/damaged DNA and senescence signals.^42,52,53^ Moreover, NF-κB and NLRP3 have both been implicated in tau and Aβ pathology,^53^ both interact with AD risk genes/proteins, ^40,42,44,46–53^ and several studies have shown that inhibiting NF-κB or NLRP3 pharmacologically or genetically protects against neurodegeneration.^40,53–55^

Inflammasome inhibition has been intensely investigated as a novel target, as well as a common cure for a range of neurodegenerative and autoimmune diseases,^56,57^ and using NF-κB and NLRP3 mRNA targeting and translation-inhibiting Nanoligomer combination (NI112) is a safe, effective, and targeted approach.^58–65^ However, even other inflammasome-targeting small-molecule therapeutics (e.g., MCC950) are limited by: 1) potential off-target effects;^66–68^ and 2) the fact that they target only NF-κB or NLRP3 alone. NI112, an RNA therapeutic that crosses the blood-brain barrier, addresses problems with pan immunosuppressive therapies (such as steroids), and the cocktail potently inhibits neuroinflammation.^58–64^ Thus, whereas clinical trials of generic anti-inflammatory drugs for cognitive decline/AD have been disappointing,^69–71^ targeting neuroinflammation with NI112 has promising therapeutic potential. We have shown that NI112 reduces pro-inflammatory astrocytes/microglia even in prion-treated mice,^61^ aging-associated neuropathology and AD,^64^ and EAE mice^63^ (models of severe neurodegeneration), and that this is associated with cognitive improvements. Moreover, the high binding specificity (K_D_ ∼5 nM^60,62,72^), minimal off-targeting,^60,62,65^ high safety profile,^65^ lack of any immunogenic response or accumulation in first pass organs,^65^ absence of any observable histological damage to organs even for long (>15-20 weeks) treatments,^60–62,65^ and facile delivery to the brain to counter neuroinflammation,^58,61,63,65^ make Nanoligomers and specifically the NI112 cocktail a promising, safe and targeted approach to mitigate neuropathological inflammation, as well as microglial and astrocytic activation.

## RESULTS AND DISCUSSION

### ALS therapeutic evaluation in human 3D motor neuron (MN) organoids

We developed a high-throughput screening approach for testing ALS and AD therapeutics using human 3D motor neuron organoids (**Fig. 1A**). Using a modification of published protcols^73,74^ with 90:10 human neuron: astrocyte combination, we created 3D MN organoids (see details in Methods). The healthy induced pluripotent stem cell (iPSC) derived motor neurons were used to create healthy MN organoids (HMN) as negative/healthy controls, and a TDP-43 knock-in/overexpressing iPS MN neurons were used to create diseased ALS organoids (or ALS). Following the 3-week maturation period, we treated the MN organoids with 10 µM NI112 for 4 weeks and conducted two kinds of tests: 1) functional testing using calcium imaging; and 2) biochemical testing of cell supernatant/serum and lysed organoids (at the end of the 4-week experiment) using TDP-43 and other ALS biomarker ELISA. As outlined above, TDP-43 has been shown to be a key risk and pathological factor for ALS,^10–13^ as well as other neurodegenerative diseases.

**Fig. 1.**
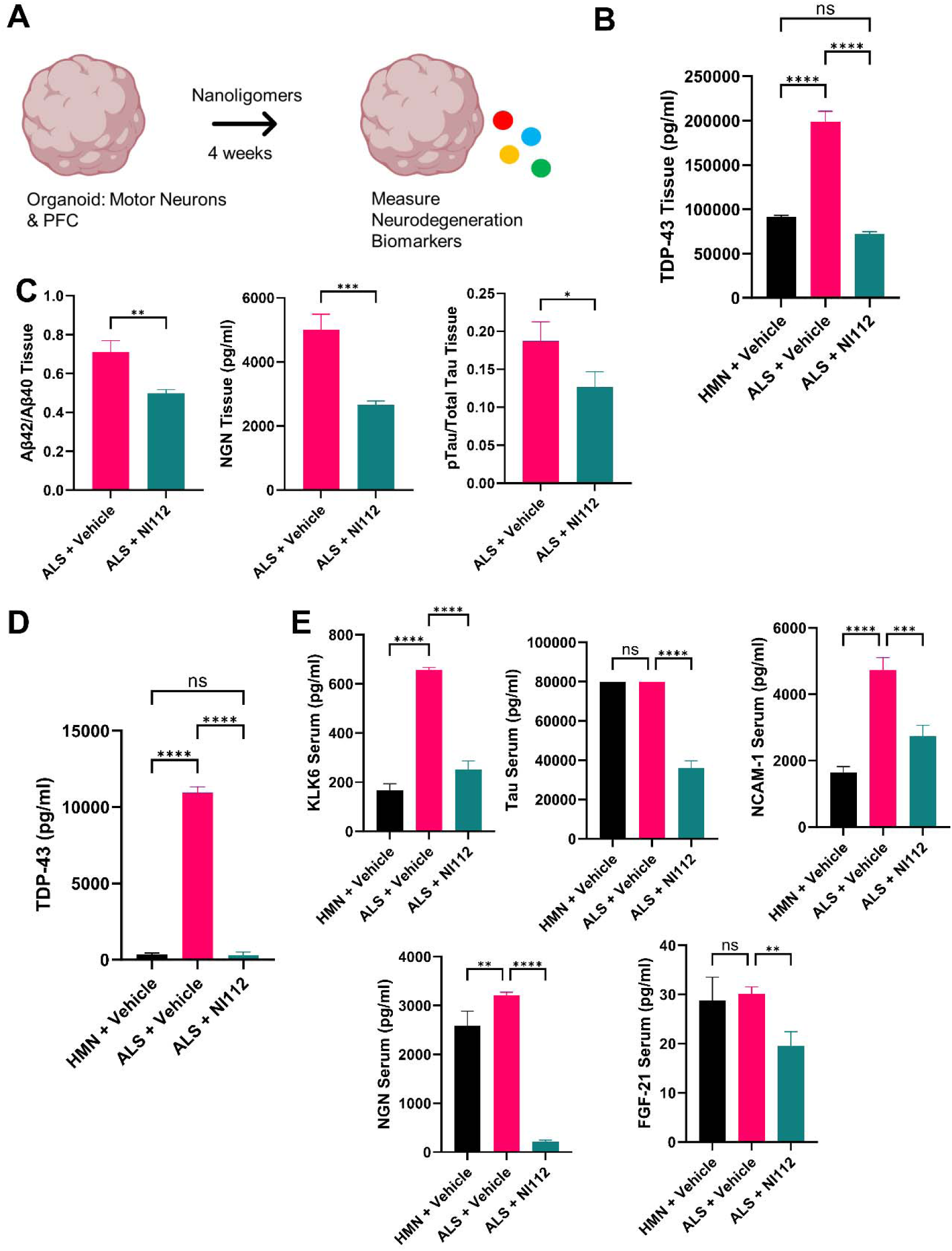
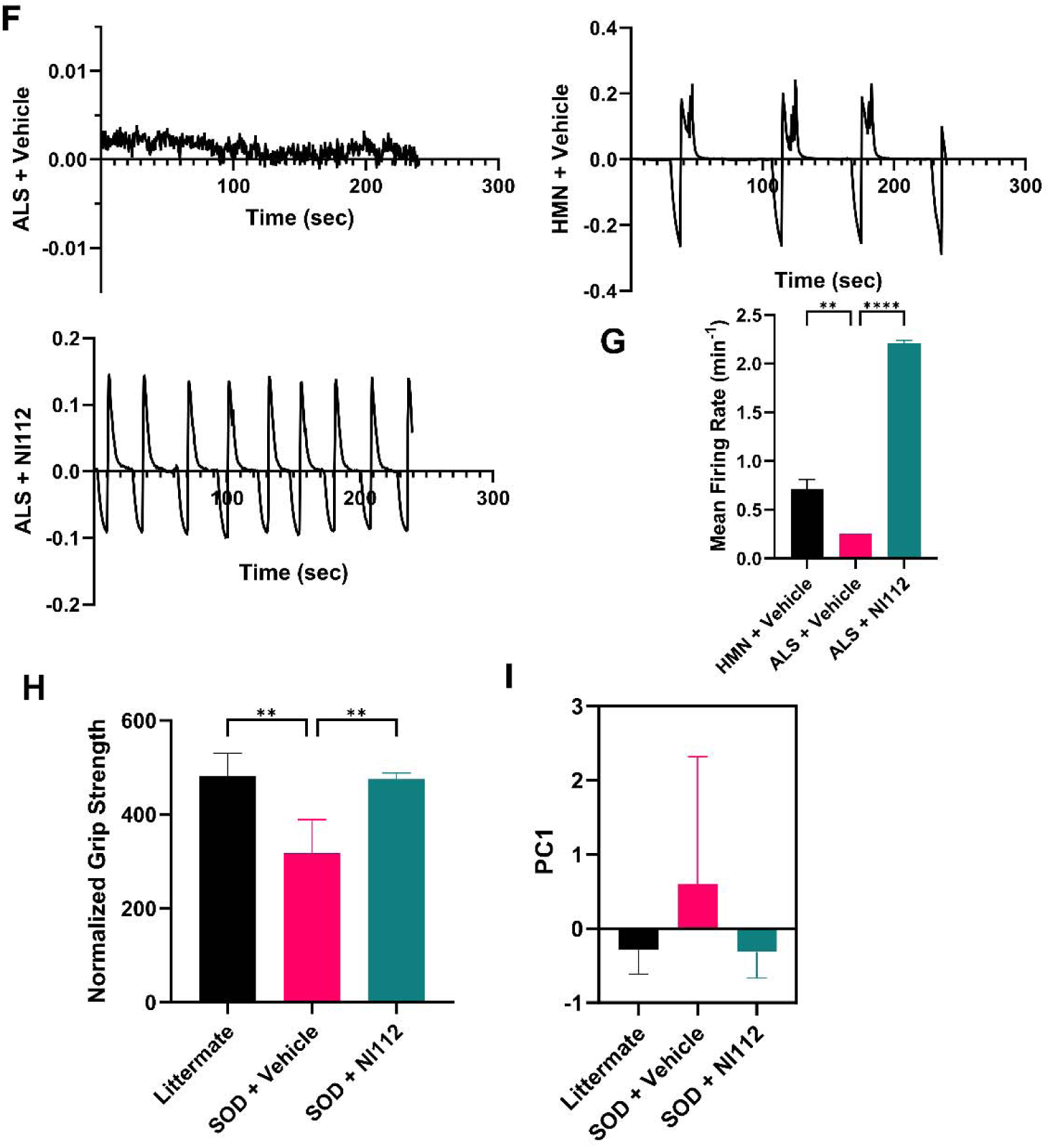
In *vitro* and *in vivo* validation of Nanoligomer NI112 treatment in healthy (HMN) and diseased (ALS) MN organoids, and SOD1-G93A ALS mouse model. **A.** Schematic showing *in vitro* treatment and testing using 4-week Nanoligomer treatment in diseased 3D human organoids. ALS MN organoids were prepared through TDP-43 overexpressing MNs, whereas AD organoids shown here had APOE4/APOE4 double allele mutation. The organoid lysate and supernatants were monitored using different neurodegenerative disease biomarkers. **B**. TDP-43 protein expression in organoid tissue for untreated healthy (HMN + Vehicle, negative control), sham-treated diseased organoids (ALS + Vehicle, positive control), and compared to Nanoligomer-treated diseased organoid (ALS + NI112) supernatant, for testing the potential efficacy of NI112 Nanoligomer treatment in the ALS disease model (using TDP-43 as a disease biomarker). **C**. Additional disease biomarkers like amyloid beta 1-42 (Aβ42), amyloid beta 1-40 (Aβ40), Neurogranin (NGN), phosphorylated tau pT181 (pTau), and Toal Tau protein (Total Tau), all demonstrate a significant reduction in ALS disease pathology in the disease MN organoids (ALS) with NI112 4-week treatment. Corresponding measurements of these biomarkers in organoid supernatants at the end of 4 weeks also confirm a reduction in disease pathology measured through: **D**. TDP-43, and **E**. other potential ALS biomarkers like Kallikrein-6 (KLK-6), Total Tau, Neural cell adhesion molecule-1 (NCAM-1), NGN, and Fibroblast growth factor 21 (FGF-21). **F.** Representative calcium signaling traces for healthy MN (HMN + Vehicle), ALS untreated MN organoids (ALS + Vehicle), and ALS NI112 treated (ALS + NI112) organoids. **G.** Averaged mean firing rate for organoids in all groups. **H.** Behavioral/functional measurement of grip strength used to assess ALS disease progression in SOD1-G93A transgenic mouse model. NI112 treatment restored the behavioral outcome with a 4-week treatment. Littermate mice (negative control group), SOD1-G93A sham-treated mice (SOD + Vehicle, positive control group), and SOD1-G93A Nanoligomer-treated mice (SOD + NI112, treatment group), were compared to assess the functional impact of the treatment on motor function (measured using rotarod setup). **I.** Principal component analysis (PC1) showing treated mice looked similar to negative control (littermate) mice, rather than untreated SOD (positive control) mice, after 4-week treatment. Inset shows a zoomed-in image of areas marked using a square. NI112 used in human organoids is a Nanoligomer targeting the NFκB gene (A-G), whereas NI112 used in mouse models is a cocktail of Nanoligomers targeting both NFκB and NLRP3 genes (H-I). \**P* < 0.05, \*\**P* < 0.01, and \*\*\**P* < 0.001, \*\*\*\**P* < 0.0001, Mean ± SEM, significance based on one-way ANOVA. *n* = 3 for each group. *In vitro* concentration: 10 µM. *In vivo* dose: 150 mg/kg; Method of administration: intraperitoneal (IP) injection.

### Reduction of neuropathology and improved neuronal firing with NI112 treatment in ALS organoids

We observed a significant reduction in ALS pathology, measured using TDP-43 and other neurodegenerative biomarker ELISAs, both in lysed organoid tissue (**Fig. 1B, C**) and cell supernatant/serum (**Fig. 1D, E**). Besides the clear reduction of TDP-43 expression in lysed tissue from ALS organoids (levels at par with the healthy MN organoids, **Fig. 1B**), an important ALS biomarker,^10–13^ we also found reductions in other neurodegeneration biomarkers like Kallikrein-6 (KLK6), Amyloid Beta 42 (Aβ42) and phosphorylated tau (pTau) in NI112-treated vs. sham-treated ALS organoids (**Fig. 1C**). KLK6 is an important neurodegenerative biomarker shown to be directly linked to axonal and neuronal degeneration and motor function loss,^75^ as well as other neuropathologies such as neuroinflammation and Tau phosphorylation leading to identification in other CNS disease biomarkers.^76^ Additionally, Aβ42^77^ and pTau (pT181),^78^ which are associated with AD, were also elevated in ALS^77,78^ organoids and reduced to their HMN levels with NI112 treatment. Additional neurodegeneration biomarkers including NCAM-1,^79^ Neurogranin (NGN), ^80,81^ and FGF-21 (**Fig. 1E**) were also found to be significantly downregulated upon NI112 treatment. These biomarkers are relevant for aging-related cognitive decline, and importantly regulate synaptic activity of neurons^79–81^ In fact, the ALS NI112-treated organoids were statistically similar to the HMN organoids, indicating potential biomarker restoration to a healthy state/mitigation of ALS pathology. Finally, when we measured the neuronal firing rates in MN organoids, we observed a significant decrease in untreated ALS organoids (positive control, **Fig. 1F, G**) mean firing rate, compared to healthy MN organoids (HMN), but the NI112-induced TDP-43 biomarker reductions were accompanied by significant improvement in mean firing rate, even when compared to HMN negative controls (**Fig. 1G**).

### NI112 Evaluation in SOD1-G93A ALS mouse model

We next assessed motor function and the effect of NI112 treatment *in vivo* in the ALS SOD1-G93A mouse model of ALS.^82–84^ Given the neuromuscular degeneration observed in ALS, we utilized established grip strength tests to detect the development of ALS-like neuromuscular dysfunction in this transgenic mouse strain. Since SOD1 is a more reproducible mouse model of ALS with predictable model performance compared to TDP-43 mouse models,^82–84^ we used Littermates (negative control), SOD + Vehicle (saline/sham-treated), and SOD + NI112 treatment groups. We observed a significant decrease in measured grip strength for mice between 11-15 weeks of age (post onset of symptoms, so after ALS diagnosis in patients), as shown in literature,^82–84^ for negative and positive control groups (**Fig. 1H**). However, NI112 treatment showed significant improvement in this behavioral output, with just 4 weeks of treatment (at 15 weeks of age).

### Biochemical assessment for NI112 treated mice

To evaluate the biochemical effects of NI112 in the mice we studied, we harvested the brain after 15 weeks of age (4 weeks of NI112 treatments) and measured inflammation in the Thalamus region for the three groups. In ALS-model mice, we observed a consistent increase in inflammatory cytokines and chemokines (**Fig. S1, 1I**), and a combined One-way ANOVA analysis for all cytokines showed significance between Littermate and SOD + Vehicle groups (p-value=0.0033) and SOD + Vehicle and SOD + NI112 groups (p-value=0.0008), but not between Littermate and SOD + NI112 group (p-value=0.5811). Further analysis using Principal component Analysis (PC1 43.4% variance) also showed Littermate and SOD + NI112 groups were similar (more negative values, showing lower inflammation, **Fig 1I**), whereas SOD + Vehicle group showed higher inflammation (more positive values), although with higher variance possibly due to small group sizes.

### Testing AD therapeutic in 3D human organoids

As in our ALS organoid studies above, we also developed a high-throughput screening method using human 3D prefrontal cortex (PFC) organoids for testing AD therapeutics (**Fig. 1A**). Using published protcols^73,74^ with 90:10 human neuron: astrocyte combination, we created healthy (HPFC) organoids (see details in Methods). Briefly, healthy glutamatergic and Gamma-Aminobutyric Acid (GABA)-ergic (70:30) neurons were used to create healthy PFC organoids (or HPFC) as negative/healthy controls, and an APOE4/APOE4 double allele mutation in the GABA-ergic neuronal cell line was used to create diseased AD organoids (or AD). Following the 3-week maturation period for PFC organoids, we treated the organoids with 10 µM NI112 for 4 weeks and tested for amyloid plaques using Aβ42 as the key biomarker,^85,86^ and other AD biomarkers using ELISA.

### Reduced Amyloid plaque and other AD biomarkers with NI112 treatment in PFC-AD organoids

Healthy PFC organoids (positive control) had significantly lower amyloid plaque than diseased AD organoids (**Fig. 2A**). However, in NI112-treated AD organoids, we observed a significant reduction in Aβ plaques, measured using Aβ42 biomarker ELISA (**Fig. 2A, E**, ∼50% reduction, p<0.01). In fact, AD-NI112-treated organoids were statistically no different from healthy PFC organoids, when comparing multiple AD and neurodegeneration biomarkers (**Fig. 2A-F**). Specifically, we used pTau and total Tau (**Fig. 2B, F**),^87^ TDP-43 (**Fig. 2C**),^13,88^ and Neurogranin (NGN, **Fig. 2D**)^81^ as AD and neurodegeneration biomarkers, and we saw significant decreases in all biomarkers with 4-week NI112 treatment of AD organoids.

**Fig. 2.**
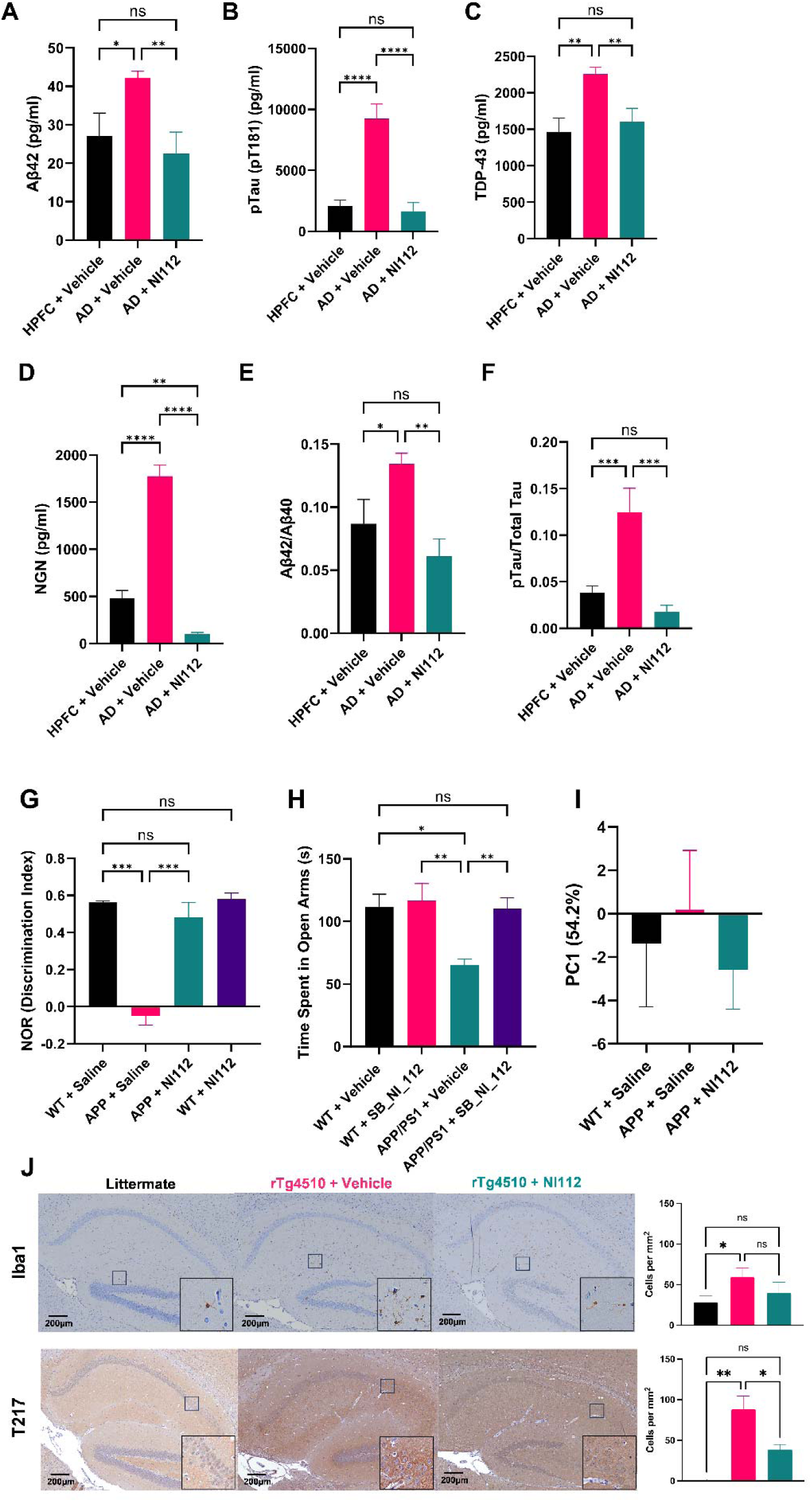
NI112 treatment is therapeutic *in vitro* (using healthy (HPFC) and diseased (AD) organoids) and *in vivo* (APP/PS1 Irradiated and Tg4510 AD mouse models). 4-week treatment of PFC organoids made from healthy (HPFC) and APOE4/APOE4 diseased organoids (AD) reveals a significant reduction in multiple AD biomarkers. Comparison between untreated healthy (HPFC + Vehicle, negative control), sham-treated AD (AD + Vehicle, positive control), and Nanoligomer treatment group (AD + NI112) measured using: **A.** Amyloid beta 1-42 (Aβ42); **B.** phosphorylated tau pT181 (pTau); **C.** TDP-43; **D.** Neurogranin (NGN); **E.** Aβ42/Aβ40; **F.** pTau/Total Tau. All AD biomarkers show significant reduction in AD organoids, at levels comparable to healthy PFC organoids. **G.** Behavioral measurement of cognition (using novel object recognition, NOR) used to assess AD disease progression in this transgenic APP/PS1 mouse model. NI112 treatment restored the behavioral outcome with a 4-week treatment. Sham-treated wild-type C57BL/6 (WT + Saline, negative control group), sham-treated APP/PS1 mice (APP + Saline, positive control group), and APP/PS1 Nanoligomer-treated mice (APP + NI112, treatment group), were compared to assess any improvement in cognition with the 4-week NI112treatment. WT mice were also treated with NI112, to demonstrate no change in cognition, indicating the improved cognition in APP/PS1 mice is the result of reduced AD pathology. All mice groups were irradiated with Fe56 radiation to accelerate the AD pathology and neurodegeneration. **H**. Behavioral Assessment using Elevated Maze Plus test. Improved time spent in open arms also demonstrates lower anxiety in APP/PS1 mice with NI112 treatment. **I.** Principal component analysis (PC1) showing treated mice looked similar to negative control (WT) mice, rather than sham-treated APP/PS1 (positive control) mice, after 4-week treatment. **J.** Immunofluorescence staining of the hippocampus using Iba1 (for microglial staining) and T217 (for phosphorylated tau staining) in littermate (negative control), vehicle-treated rTg4510 mice (positive control), and NI112 treated rTg4510 mice (treatment group). The statistical analysis of multiple histological stains shows a significant reduction in both microglial activation and phospho-tau pathology, with 4-week NI112 treatment. NI112 used in human organoids is a Nanoligomer targeting the NFκB gene (A-F), whereas NI112 used in mouse models is a cocktail of Nanoligomers targeting both NFκB and NLRP3 genes (G-J). \**P* < 0.05, \*\**P* < 0.01, and \*\*\**P* < 0.001, \*\*\*\**P* < 0.0001, Mean ± SEM, significance based on one-way ANOVA. *n* = 3 for each group, except n=5 for APP+NI112 group and n=6 for rTg4510 mice groups. *In vitro* concentration: 10 µM. *In vivo* dose: 150 mg/kg; Method of administration: intraperitoneal (IP) injection.

### NI112 Evaluation in Irradiated APP/PS1 mouse model

Based on the above results and similar to our ALS model studies, we next tested the efficacy of NI112 *in vivo* using two separate AD models: 1) APP^swe^, PSEN1^dE9^ (APP/PS1) mice as an amyloid plaque model,^89^ and 2) rTg(tau^P301L^)4510 mice as a tau pathology model.^90,91^ To accelerate the neuropathogenesis in APP/PS1 mice, we used Fe^56^ irradiation to increase protein misfolding and amyloid pathogenesis rate (see Methods).^92^ Similar to previous reports on cognitive function (measured using novel object recognition, NOR) in these models, 3-4-month-old APP/PS1 mice showed a marked decrease in NOR 6 weeks post-irradiation. However, while age-matched wild-type (WT) mice showed no change, APP/PS1 mice showed improvement in cognitive performance with NI112 treatment post-irradiation (**Fig. 2G, H**). Further, evaluation of hippocampal brain tissue showed a marked increase in sham-treated APP/PS1 mice (APP+ Saline, positive control) compared to WT+ Saline (negative control) mice (**Fig. S2**). The 6-week NI112 treatment also strongly reduced neuroinflammation, especially in key immune cell infiltration and mobilization cytokines like IL-17 (**Fig. S2A**), and activators of immune responses like IL-23 (**Fig. S2B**). In fact, a combined principal component analysis (PC1 54.2% variance) showed WT+ Saline (negative control) mice and APP + NI112 groups were similar (more negative values, showing lower inflammation), whereas the APP + Vehicle group (positive control) had higher overall inflammation (more positive values, **Fig. 2I**). Moreover, whereas there was a clear difference between negative and positive controls (WT+ Saline and APP+ Saline, p=0.0239) and treatment groups compared to the positive control (APP+ Saline and APP+ NI112, p=0.004), there was no significance between negative control and treatment group (p=0.7547), suggesting a normalization of neuroinflammation in these treated animals. We observed similar results in terms of cognitive improvements and neuroinflammation reductions in the rTg4510 AD mouse model with NI112 4-week treatments.^64^ Further neuropathological assessment using immunohistochemistry also showed a clear effect of NI112 treatment in the rTg4510 AD mouse model (**Fig. 2J, S3**).

### Immunohistochemistry assessment of NI112 treatment in rTg4510 AD mouse model

Last, we evaluated 4-week NI112 treatment in rTg4510 tau AD model.^90,91^ Using immunohistochemical staining of activated microglia (Iba1+) as the causal neuroinflammation mechanism, and phospho-Tau AD pathology (Phospho-T217+) of the hippocampus (**Fig. 2J**) and cortex tissue (**Fig. S3**), we observed a statistically significant reduction in both neuropathologies in Tg4510 mice. Our assessment showed a clear effect of NI112 treatment leading to reduced AD disease progression in mice brains in this *in vivo* disease model.

## CONCLUSIONS

In summary, the NF-κB1- and NLRP3-inhibiting Nanoligomer cocktail NI112 can be therapeutic in: 1) ALS (TDP-43 overexpressing MN organoid) and AD (APOE4/APOE4 allele double mutation) 3D human organoid models; and 2) *in vivo* in SOD1-G93A (ALS), APP/PS1 and Tg4510 (AD) mouse models. The Nanoligomer NI112 effectively reduced ALS biomarkers in human MN organoids such as TDP-43, reduced neuroinflammation, and effectively restored motor function in *in vivo* SOD1-G93A mice (measured in rotarod tests as grip strength). Similarly, this neuroinflammation-targeting Nanoligomer cocktail improved cognition and reduced AD-related pathology in APP/PS1 (as accelerated amyloid plaque model) and rTg4510 (as Tau model) mice. These outcomes were comparable to similar measures in negative control littermate mice (without ALS) and WT/littermate mice (without AD), significantly reducing neuropathology in both studies. Together, these studies show the potential for further testing and development of these novel, neuroinflammation-targeting Nanoligomers as a new therapeutic approach in ALS and AD, with the capacity to address various neurodegenerative diseases and potentially benefit patients suffering from debilitating neuropathological conditions.

## Supporting Information

Supporting figures S1-S3, and Methods and Materials.

## Author Contributions

Conceptualization: P.N.

Data curation: S.S., D.W., S.R., V.S.G., A.C., J.A.M., T.J.L. P.N.

Formal analysis: P.N., T.J.L., A.C., J.AM., D.W., S.R.

Funding acquisition: P.N., A.C., J.A.M., T.J.L.

Investigation: S.S., D.W., S.R., V.S.G., A.C., J.A.M., T.J.L. P.N. Methodology: P.N., A.C., J.A.M., T.J.L.

Project administration: P.N., A.C., J.A.M., T.J.L.

Roles/Writing - original draft: P.N.

and Writing - review & editing: P.N., T.J.L., S.S., D.W., S.R., V.S.G., A.C., J.A.M.,

## ACKNOWLEDGMENTS

Authors acknowledge financial support from NASA SBIR Awards 80NSSC22CA116 and 80NSSC23CA171, and Dr. Michael Weil for help with Fe^56^ mice irradiation at Brookhaven National Lab.

## MATERIALS AND METHODS

### Nanoligomer design and synthesis

Nanoligomers were specifically designed and synthesized (Sachi Bio) following established methods detailed in published references.^58–62^ The Nanoligomers are composed of an antisense peptide nucleic acid (PNA)^58,59^ conjugated to a gold nanoparticle.^58,59^ The 17-base-long PNA sequence was selected to optimize solubility and specificity, targeting the start codon regions of *nfkb1* (Sequence: AGTGGTACCGTCTGCTA) and *nlrp3* (Sequence: CTTCTACTGCTCACAGG), within the Mouse genome (GCF_000001635.27), identified using our bioinformatics toolbox. Various PNA sequences were screened for their solubility, self-complementing sequences, and potential off-target interactions within the mouse genome (GCF_000001635.27). The PNA portion of the Nanoligomer was synthesized on a Vantage peptide synthesizer (AAPPTec, LLC) employing solid-phase Fmoc chemistry. Fmoc-PNA monomers, wherein A, C, and G monomers were shielded with Bhoc groups, were obtained from PolyOrg Inc. Post-synthesis, the peptides were attached to gold nanoparticles and subsequently purified using size-exclusion filtration. The conjugation process and the concentration of the refined solution were monitored by measuring absorbance at 260 nm (for PNA detection) and 400 nm (for nanoparticle quantification).

### Culturing and treating 3D human organoids with Nanoligomers

We slightly modified the published protocols and methods to culture 3D organoids.^73,74^ Briefly, both the diseased (ALS and AD), as well as healthy MN and PFC models consisted of a 90:10 Neuron: Astrocytes ratio. Cells were seeded in ultralow attachment (Sbio) plates and centrifuged at 1000 x g for 5 min. A 50% media exchange was performed every 2-3 days for 3 weeks using either Complete Motor media for MN models or Complete BrainPhys Media for PFC models until organoids reach maturation. For the first week of growth, MN organoids were fed using Complete Motor Media with a concentration of 5µM DAPT (Cayman Chemical). Post 7 days of seeding, they were fed with Complete Motor Media without DAPT per the user’s manual instructions. Typical PFC organoids used 90:10 Neuron: Astrocyte ratio with 70:30 distribution of glutamatergic: GABAergic neurons. On day 21, organoids were treated with 10µM (effective concentration) Nanoligomer (NI112). Relevant media was added to No Treatment Controls. Following 24-hour treatment, the supernatants were collected and analyzed using multiplexed ELISA per manufacturer recommendations, as detailed below.

### Organoid calcium staining and dynamics

After the completion of the treatment, organoids were rinsed twice with the corresponding serum-free media and then treated with Calbryte™ 520 (AAT Bioquest, 10µM effective concentration) prepared in corresponding serum-free media. Organoids were incubated at 37°C, 5.0% CO_2_ for three hours, rinsed twice with the corresponding complete media, and imaged for calcium dynamics using a Nikon Widefield fluorescent microscope in a humidified chamber maintained at 37°C and 5% CO_2_.

### Organoid lysis and protein extraction

Three organoids were combined in a 1.5 mL Eppendorf tube and washed with DPBS (Gibco). Cells were first dissociated by adding TrypLE (Gibco) and incubated at 37°C 5.0% CO2. After that, appropriate complete (Brain Phys or Motor) media was added to neutralize TrypLE, cells were centrifuged, and washed twice with DPBS. Following this, DPBS was completely aspirated, and Bio-Plex Cell-Lysis Kit (Bio-Rad) was added. Following Bio-Rad user’s instructions for cell lysis, the organoids were sonicated, centrifuged at high speed, and the supernatants of the lysates were collected for analysis of secreted protein/cytokine expression using multiplexed ELISA.

### In vivo testing in SOD1-G93A ALS mice model

8-9-week-old female C57Bl/6SOD1-G93A mice and corresponding littermate controls were purchased from Jackson Labs. Nanoligomer cocktail (1x intraperitoneal injection; 150 mg/kg body weight) or vehicle was started on Day 2 week 11, and then given 3 until week 15. Grip strength was tested in mice using the DST-110 Digital Force Gauge (Maze Engineers). 5 trials were performed per mouse and the highest and lowest values were discarded for analysis. The remaining 3 values were averaged for each mouse. Data are presented as g force/g body weight as previously described. Mice were housed throughout all experiments at ∼18-23°C on a 12 light/12 dark cycle. Fresh water and *ad-libitum* food (Tekland 2918; 18% protein) were routinely provided to all cages. Animals were consistently health checked by the veterinary staff at Colorado State University (CSU). This protocol was approved by CSU IACUC and Laboratory Animal Resources staff.

### In vivo testing in irradiated APP/PS1 AD mice model

8-9-week-old female APP/PS1 and WT C57Bl/6 mice were purchased from Jackson Labs. The mice were shipped to Brookhaven National Lab and irradiated with 1Gy Fe56 radiation for accelerated neurodegeneration mimicking AD pathology.^92^ The mice were then shipped back to the CSU vet core and their behavioral assessment started. 1-week post-irradiation, NI112 Nanoligomer cocktail (1x intraperitoneal injection; 150 mg/kg body weight) or vehicle was started, and then given until 6-8 weeks after irradiation, after which the mice were euthanized for biochemical and immunofluorescence assessment. Mice were housed throughout all experiments at ∼18-23°C on a 12 light/12 dark cycle. Fresh water and *ad-libitum* food (Tekland 2918; 18% protein) were routinely provided to all cages. Animals were consistently health checked by the veterinary staff at Colorado State University (CSU). This protocol was approved by CSU IACUC and Laboratory Animal Resources staff.

### In vivo testing in rTg4510 AD mice model

8-9-week-old 3 male and 3 female rTg4510 mice were purchased from Jackson Labs, along with corresponding littermate controls. Nanoligomer cocktail (1x intraperitoneal injection; 150 mg/kg body weight) or vehicle was started after their 1-week acclimatization, and then given until 4 weeks after which the mice were euthanized and assessed for Ad pathology using immunofluorescence. Mice were housed throughout all experiments at ∼18-23°C on a 12 light/12 dark cycle. Fresh water and *ad-libitum* food (Tekland 2918; 18% protein) were routinely provided to all cages. Animals were consistently health checked by the veterinary staff at Colorado State University (CSU). This protocol was approved by CSU IACUC and Laboratory Animal Resources staff.

### Tissue collection

After deep anesthetizing with isoflurane, ∼1 ml of blood was removed via cardiac puncture followed by cervical dislocation. The left hippocampus and a piece of left cortex tissue were removed, flash-frozen on dry ice, and stored at −80°C until further processing. The right hippocampus and right cortex were processed (paraffin-embedded) for immunostaining (details below). The spinal cords were also collected with the cervical and lumbar regions preserved in 10% normal buffered formalin (NBF) and the thoracic portion was flash-frozen on dry ice, and stored at −80°C until further processing.

### Multiplexed ELISAs

Organoid culture supernatants and lysates were analyzed for secreted proteins using ProcartaPlex™ Human Neurodegeneration Panel 1, 9-plex (Thermofisher Scientific, Carlsbad, CA) Tissue homogenates from all mice underwent evaluation utilizing the ThermoFisher Procartaplex cytokine/chemokine panel, following previously documented procedures.^58,59^ Briefly, 25µL of 10 mg/ml brain tissues were homogenized using ProcartaPlex™ Cell Lysis Buffer (ThermoFisher) using 5-mm stainless steel beads (Qiagen) at 25 Hz for 1.5-3min (Qiagen/Retsch Bead Beater). Following homogenization samples were centrifuged at 16,000 x g for 10 mins at 4 °C. After centrifugation, homogenized samples were measured for protein concentration using DC Protein Assay (Bio-Rad) on a Tecan GENios microplate reader.

Mice brain homogenates, organoid lysates, and supernatants were then assessed in accordance with the manufacturer’s provided protocol and analyzed using a Luminex MAGPIX xMAP instrument. Standards for each cytokine/chemokine were employed at 1:4 dilutions (8-fold dilutions), alongside background and controls. Subsequently, concentrations of samples were determined from a standard curve using the Five Parameter Logistic (5PL) curve fit/quantification method.

### Immunohistochemistry

Paraffin-embedded mice brains were sectioned at 4 µm before staining, and then deparaffinized, rehydrated, sodium citrate treated, and blocked in Tris A/2% donkey serum (Jackson Immuno Research), then incubated overnight in the following IBA1 antibody (1:50 dilution; Abcam). Wash steps were then performed using 2% bovine serum albumin and 2% Triton-X in 1LM TBS, and an ABC HRP peroxidase detection kit (Vector Laboratories) and ImmPACT DAB Substrate Peroxidase (HRP) Kit (Vector Laboratories) were used for a chromogen. Slides were counterstained with hematoxylin (ThermoFisher), secured with a coverslip in mounting medium, and stored at room temperature until imaging. Whole tissue images were taken on an Olympus BX53 microscope with an Olympus DP70 camera using an Olympus UPlanSApo 20x objective (N.A. = 0.75). Representative images were taken using an Olympus BX53 microscope with an Olympus DP70 camera using an Olympus UPlanFL N 40x objective (N.A. = 0.75).

---

**Fig. S1.**
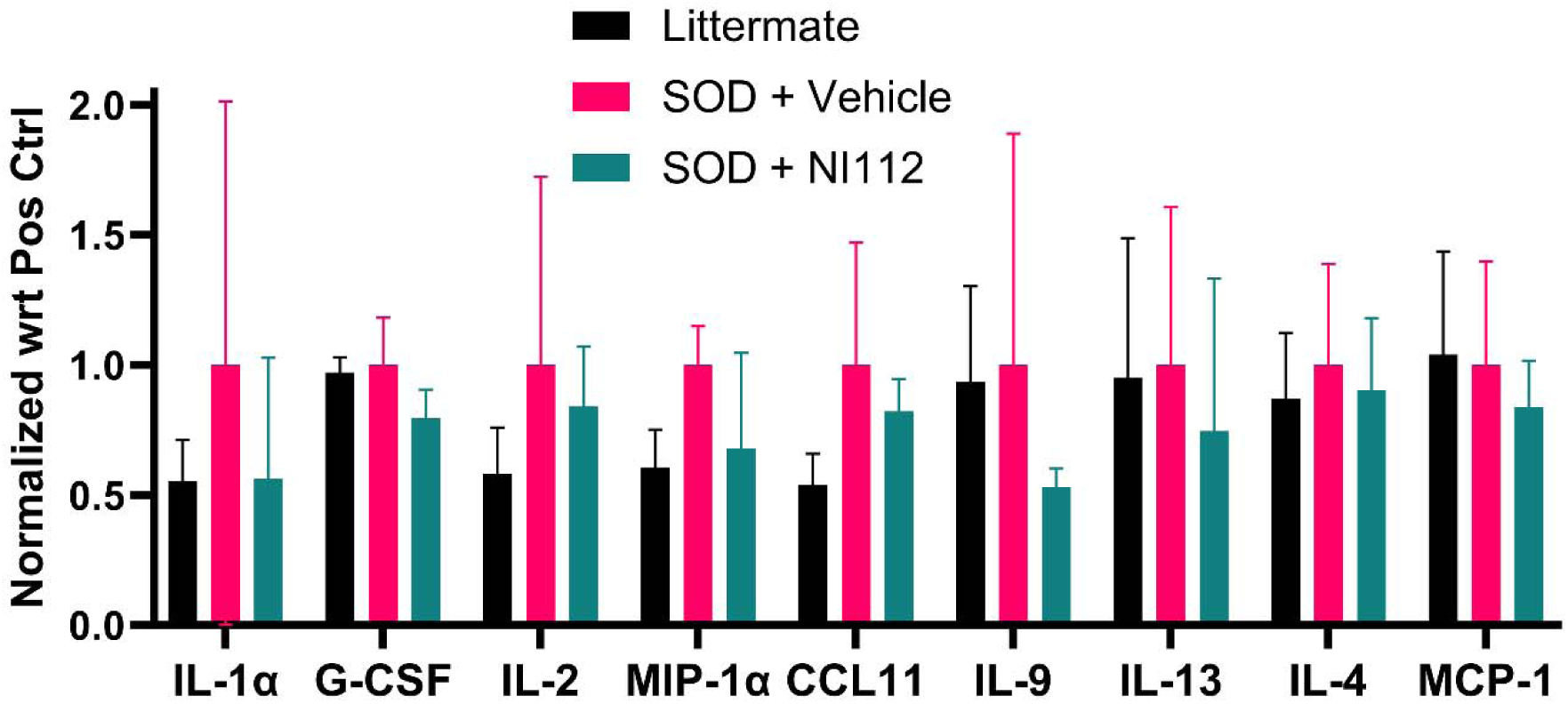
NI112 treatment is therapeutic *in vivo* SOD1-G93A ALS mouse model. Measurement of inflammatory cytokines in thalamus brain tissue of mice from different groups showing treated mice looked similar to negative control (littermate) mice, rather than untreated SOD (positive control) mice, after 4-week treatment. \**P* < 0.05, \*\**P* < 0.01, and \*\*\**P* < 0.001, Mean ± SEM, significance based on one-way ANOVA. *n* = 3 for each group. Dose: 150 mg/kg. Method of administration: intraperitoneal (IP) injection.

**Fig. S2.**
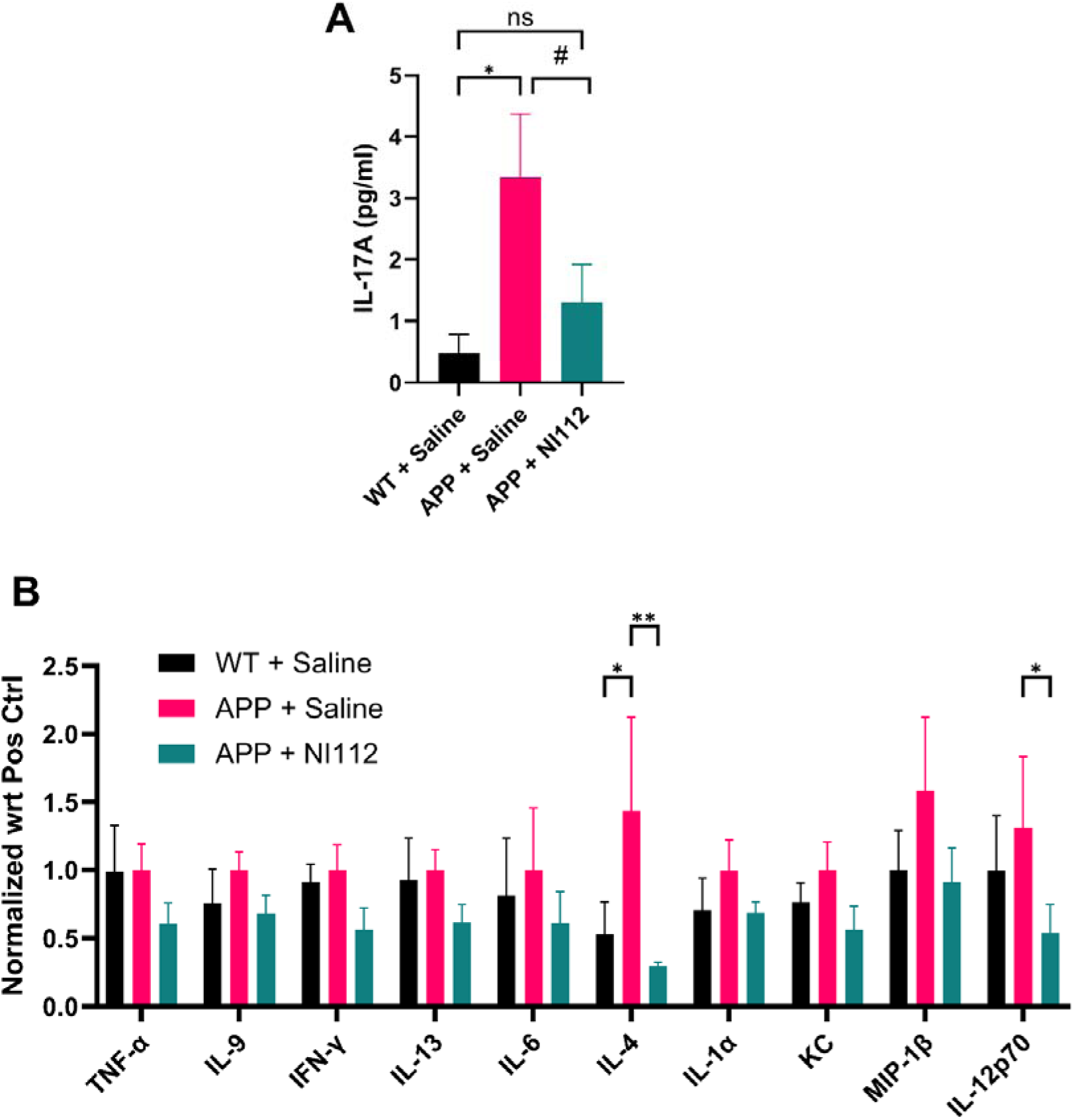
NI112 treatment is therapeutic *in vivo* APP/PS1 Irradiated AD mouse model. Measurement of inflammatory cytokines in hippocampus brain tissue of mice from different groups. Sham-treated wild-type C57BL/6 (WT + Saline, negative control group), sham-treated APP/PS1 mice (APP + Saline, positive control group), and APP/PS1 Nanoligomer-treated mice (APP + NI112, treatment group), were compared to assess any change in inflammatory cytokines with NI112treatment. **A.** IL-17, and **B.** TNF-α, IL-9, IFN-γ, IL-13, IL-6, IL-4, IL-1 α, KC, MIP-1β, and IL-12p70. \**P* < 0.05, \*\**P* < 0.01, and \*\*\**P* < 0.001, Mean ± SEM, significance based on one-way ANOVA. *n* = 3-5 for each group (n=3 for WT + Saline, WT + NI112, and APP + Saline groups, n=5 for APP + NI112 group). Dose: 150 mg/kg. Method of administration: intraperitoneal (IP) injection.

**Fig. S3.**
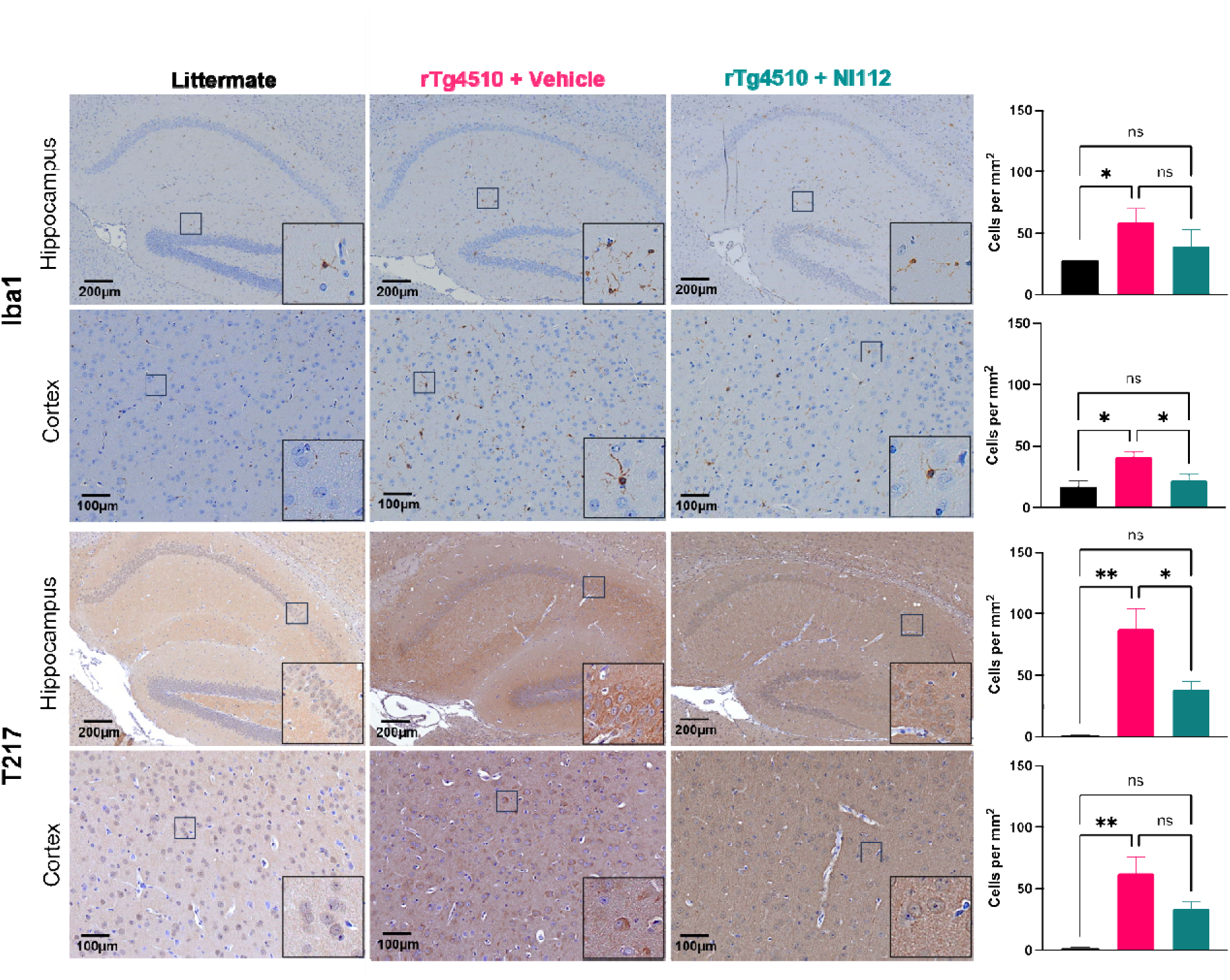
NI112 treatment is therapeutic *in vivo* rTg4510 AD mouse model. Immunofluorescence analysis of hippocampus and cortex brain tissue of mice from different groups. Sham-treated littermate mice (Littermate, negative control group), sham-treated rTg4510 mice (rTg4510 + Vehicle, positive control group), and Nanoligomer-treated rTg4510 mic (rTg4510 + NI112, treatment group), were compared to assess any change in microglial activation (with Iba1 staining), and tau pathology (phosphor-tau staining with T217) with NI112treatment. The plots on the right show combined analysis from several images showing clear statistical differences between the controls and the treatment group. \**P* < 0.05, \*\**P* < 0.01, and \*\*\**P* < 0.001, Mean ± SEM, significance based on one-way ANOVA. *n* = 6 for each group. Dose: 150 mg/kg. Method of administration: intraperitoneal (IP) injection.

